# Regulation of transcription termination by FUS and TDP-43

**DOI:** 10.1101/788778

**Authors:** Dorothy Yanling Zhao, Zuyao Ni, Shuye Pu, Guoqing Zhong, Frank W. Schmitges, Ulrich Braunschweig, Benjamin J. Blencowe, Jack F. Greenblatt

## Abstract

The carboxy-terminal domain (CTD) of the RNA polymerase II (RNAPII) subunit POLR2A is a platform for modifications specifying the recruitment of factors that regulate transcription, mRNA processing, and chromatin remodelling. We previously found that symmetrical dimethylation (me2s) of a CTD Arginine residue (R1810 in human) causes recruitment of the Tudor domain of SMN, which interacts with Senataxin. SMN is mutated in spinal muscular atrophy (SMA), and Senataxin is sometimes mutated in Amyotrophic Lateral Sclerosis (ALS). R1810me2s and SMN, like Senataxin, are important for resolving R-loops (DNA:RNA hybrids) at transcription terminators. FUS and TDP-43 (TARDBP) are DNA/RNA binding proteins that are sometimes mutated in ALS and FTD (Frontotemporal dementia). Here we show that TDP-43 and, to some extent, FUS are recruited by the R1810me2s-SMN pathway. Defects in FUS and TDP-43 recruitment influence RNAPII termination and R-loop accumulation, leading to elevated DNA damage at terminators that may contribute to neurodegenerative disorders like ALS and FTD.

## INTRODUCTION

The mammalian POLR2A CTD contains 52 repeats, with the N-terminal half containing mostly consensus heptads (Tyr_1_Ser_2_Pro_3_Thr_4_Ser_5_Pro_6_Ser_7_) and the C-terminal half containing many heterogeneous repeats (1). The CTD repeats can be phosphorylated on Tyr1, Thr4, Ser2, Ser5, and Ser7 residues. Ser5 phosphorylation by the general initiation factor TFIIH correlates with transcription initiation and mRNA capping (2,3). Ser2 phosphorylation by P-TEFb is important for RNAPII promoter release, elongation, and interactions with enzymes needed for mRNA 3’-end formation, termination, and histone H3K36 methylation (1,4,5). Tyr1 phosphorylation in yeast stimulates association of the elongation factor Spt6 and impairs recruitment of the termination factors Nrd1, Pcf11, and Rtt103 (6), and Thr4 phosphorylation is required for histone mRNA 3’-end processing (7). CTD R1810 is known to be asymmetrically dimethylated (Rme2a) by CARM1 (8), symmetrically dimethylated (Rme2s) by PRMT5 (9), and citrullinated by PADI2 (10).

We showed previously that the CTD R1810me2s modification leads to the recruitment of the adaptor protein SMN through its Tudor domain (9). SMN can self-associate through its N- and C-terminal domains to form a multimeric adaptor for the assembly of Rme2s-containing proteins into complexes (11,12). Because SMN participates in spliceosome assembly and is also present in cytosolic ribonucleoprotein complexes, mis-regulation in these two roles has been proposed to contribute to SMA (11). Complete loss of SMN is embryonic lethal in mammals and, in particular, mutations in SMN’s oligomerization or Tudor domain cause the SMA phenotype (13). One SMN interactor is known to be Senataxin (SETX) (14), a DNA:RNA helicase that is sometimes mutated in ALS, and both proteins participate in the transcription termination by RNAPII (9,15). Among the ∼150 human proteins that have been shown by mass spectrometry to contain dimethylarginine, a remarkable number (>50) are involved in RNA metabolism, transcription elongation and termination by RNAPII (Supplementary Table 1) (16-20). Therefore, it seemed possible that the CTD R1810me2s-SMN pathway would enhance the assembly of other Rme2s-containing proteins onto the RNAPII CTD for the proper regulation of transcriptional events.

The mutations implicated in ALS are highly heterogeneous in character (Supplementary Table 2) (21) and include an abnormal hexanucleotide expansion in C9orf72, as well as mutations in a variety of unrelated proteins, such as the mitochondrial superoxide dismutase SOD1, a secreted ribonuclease ANG, and the RNA binding proteins (RBPs) FUS and TDP-43 (22-25) that are involved in transcription regulation and alternative splicing (26,27). The C9ORF72 hexanucleotide expansion is found in ∼40% of ALS cases, and the expanded sequences are translated into dipeptide repeats that lead to the formation of inclusion bodies, which trap RBPs such as TDP-43 (28,29). The pathology caused by SOD1 mutations is also due to the formation of inclusion bodies rather than the lack of dismutase activity (30). FUS and TDP-43 are known to interact with each other and are localized in the nucleus and cytoplasm; in many types of ALS/FTD, TDP-43 or FUS is known to be trapped in cytoplasmic inclusion bodies (31,32).

Although FUS is mostly nuclear, most FUS mutations that lead to ALS are in its nuclear localization signal sequence and cause it to be trapped in cytoplasmic inclusions (33). TDP-43, also mainly a nuclear protein, is found in cytoplasmic inclusions in almost all cases of ALS (24,25) even though ALS-causing TDP-43 mutations are rare. Therefore, one hypothesis to explain ALS would be that the formation of abnormal cytoplasmic inclusions caused by mutations in many different proteins traps certain RBPs such as FUS and TDP-43, depleting them from the nucleus where they are needed for gene regulation. Among the SMN-interacting proteins, we found that TDP-43 and, to some extent, FUS act downstream of the RNAPII CTD R1810me2s-SMN pathway to resolve R-loops at terminators. Our study supports the idea that an underlying mechanistic basis for neurodegenerative disorders such as SMA, ALS, and FTD lies in the R-loop accumulation and consequent DNA damage caused by defects in RNAPII termination (9).

## MATERIAL AND METHODS

### Cell Culture, shRNA knock-downs, siRNA knock-downs, CRISPR/Cas9 knockouts, and GFP over-expressions

There was no evidence of mycoplasma contamination of the cell lines used in this work as judged by staining of fixed cells with DAPI. Raji cells were cultured in RPMI (SLRI media facility, University of Toronto) plus 10% Tetracycline free FBS (Gibco) and 1% Glutamate, and stably transduced cells were maintained with 500μg/ml G418 (Gibco, 11811031). HEK293 cells were grown in DMEM (SLRI media facility) plus FBS (Sigma, F1051), and stably shRNA knocked down transduced cell lines were maintained with 2μg/ml puromycin (Sigma, p8833). Flp-In 293 T-REx cell lines were obtained from Life Technologies (R780-07), and stable Flp-In 293 T-REx cell lines after the incorporation of the transgene were maintained with hygromycin (Life Technologies, 10687010) at 2ug/ml.

shRNAs in lentivirus vectors were used to stably transduce cell lines using an established protocol (34). siRNA knock-downs for HEK293 cells were performed with 50nM siRNAs with PepMute siRNA transfection reagent (SigmaGen Laboratory, SL100566) for 3 days. siRNAs against human FUS (NM_001010850) and TDP-43 (NM_007375) and scrambled control were purchased from Sigma.

For CRISPR/Cas9-mediated gene knock-outs, CRISPR/Cas9 plasmids (pCMV-Cas9-GFP) were purchased from Sigma-Aldrich to express the scrambled guide RNA, or guide RNAs for the KO of TDP-43 or FUS. 2 µg of the plasmids were transfected into HEK293 cells, and 1 day after lipofectamine 3000 transfection, cells were sorted by BD FACSAria flow cytometry (Donnelly Centre, University of Toronto) and single GFP-positive cells were plated and expanded into 48-well plate. The expression level of TDP-43 or FUS in each clone was detected by western blotting.

For CRISPR/Cas9-mediated knock-in mutation of POLR2A R1810 to alanine, CRISPR/Cas9 plasmids (pCMV-Cas9-GFP) express the scrambled guide RNA, or guide RNAs targeting the 29^th^ exon of POLR2A with the following sequences: 5’-CACCGGAGACTGTGGTGTGTATCGT-3’; 5’-AAACACGATACACACCACAGTCTCC-3’. We synthesized a repair DNA template of 750 base pairs (Life Technologies) that contained the desired mutation of CGA → GCA to create the R1810 → A mutation. 1 µg of the CRISPR/Cas9 plasmids and 3ug of the repair template were transfected into HEK293 cells, and 2 days after lipofectamine 3000 transfection, cells were sorted by BD FACSAria flow cytometry and single cells were plated and expanded into 96-well plates. The R1810A mutation was verified by sequencing cDNA extracted from cells derived from the individual clones.

For the stable over-expression of the TDP-43-GFP in HEK293 cells, wildtype TDP-43 ORF (Harvard Plasmid database, HsCD00079870) was cloned into the pDEST pcDNA5/FRT/TO-eGFP through Gateway cloning (ThermoFisher). The truncation mutant (TDP-43-GFP RRM-) lacking the RRM coding (S104-P262) residues was generated from the wildtype ORF template through PCR splicing/overlap extension PCR; which employs 2 complementary primers that comprise a fused sequence of −15bp to +15bp related to the junction point (5’-GCAGTCCAGAAAACA-3’+5’-AAGCACAATAGCAAT-3’) in multiple PCR reactions with a high fidelity Taq DNA polymerase (ThermoFisher, 11304011). The truncated ORF was then cloned into the pDEST pcDNA5/FRT/TO-eGFP through Gateway cloning.

### Cell transfection and electroporation

Raji cells with stable expression of HA-tagged wild type or R1810A POLR2A constructs were generated by electroporation (10μg of plasmid DNA per 10^7^ cells) followed by selection and maintenance in G418 (0.5mg/ml). α-amanitin treatment was carried out with 2μg/ml α-amanitin for 3 days for Co-IP and ChIP experiments involving HA-tagged wild type or R1810A POLR2A.

The transfections of the CRISPR/Cas9 plasmids (pCMV-Cas9-GFP) and GFP-tagged (pDEST pcDNA5/FRT/TO-eGFP) transgene into the Flp-In™ 293 T-REx cell lines were performed with FuGENE Transfection Reagent (Roche, E269A).

### Immuno-staining

For immuno-staining, cells were fixed with 4% paraformaldehyde (Sigma, P6148) and washed with PBS + 0.1% Triton × 100 (Sigma, T8532). Primary antibodies 1:50-1:100 in PBST + 30mg/ml BSA (Roche, 10735108001) were added to cells for staining overnight at 4 °C. Cells were washed with PBST and stained with Alexa Fluor 488 or 594 (Invitrogen) at 1:1000 and Hoechst 33342 (Thermo Scientific) in PBST with 5% goat serum (Sigma, G9023) at room temperature for 1 hour, followed by washes with PBS + 0.2% Triton and mounting with ImmunoMount (GeneTex, GTX30928). Images were taken using an Olympus Upright Microscope BX61 with Optigrid function and processed using Volocity OpenLab Software (PerkinElmer).

### Immunoprecipitation (IP) and western blots

IP was performed with RIPA buffer (140mM NaCl, 10mM Tris pH7.6-8, 1% Triton, 0.1% Na deoxycholate, 1mM EDTA) containing protease inhibitors (Roche, 05892791001) and Benzonase (Sigma, E1014). 1-2×10^7^ cells were lysed on ice for 25 minutes by vortexing and forcing them through a 27-gauge needle. After centrifuging at 13,000rpm for 10 minutes at 4 °C, the supernatant was incubated with 25μl (1:10 dilution) protein G beads (Invitrogen, 10-1243, 10003D) and 1-2μg of antibodies for 4 hours to overnight. The samples were washed 3 times with RIPA buffer and boiled in SDS gel sample buffer. To detect R1810me2s or R1810me2a modifications on POLR2A, alkaline phosphatase (Roche, 10108138001) treatment (5μl) at 37 °C for 30 minutes was performed for POLR2A IP samples before boiling. Samples were run using 7.5-10% SDS-PAGE and transferred to PVDF membranes (Bio-Rad, 162-0177) using a Trans-Blot^®^ SD Semi-Dry Electrophoretic Transfer Cell (BioRad, 170-3940). Primary antibodies were used at 1:250 to 1:1000 dilutions for incubation overnight, and horseradish peroxidase-conjugated goat anti-mouse, -rabbit, or –rat secondary antibodies were used at 1:10,000 (Dako, P0450). Blots were developed using SuperSignal West Pico or Femto (Thermo Scientific, 34079, 34094). Blots were quantified using ImageJ software.

### Chromatin immunoprecipitation (ChIP) and R-loop detection

ChIP was performed using the EZ-ChIP™ A - Chromatin Immunoprecipitation Kit (Millipore, 17-371) or similar homemade solutions according to the manufacturer’s instructions. Antibodies were used in the 1-2μg range, and IgG was used as a background control. R-loop detection was performed according to El Hage et al with minor modifications (35). R-loop detection was performed following the ChIP protocol except that, after the nuclear lysis and sonication, genomic DNA was de-crosslinked in ChIP elution buffer containing 5M NaCl at 65°C overnight. DNA was purified with the Qiaex II kit (Qiagen, 20021) for PCR product purification and eluted in water. S9.6 binding was carried out overnight with 25μl of Dynabeads® protein G beads (Invitrogen, 100-03D) and 1μg of antibody purified from the S9.6 hybridoma cell line (36) that recognizes RNA/DNA hybrids. Immunoprecipitated and input DNAs were used as templates for qPCR. RNase-sensitivity analysis for R-loop was carried out by adding 50 U of RNase H (Invitrogen, 18021-014) in 10X RNase H buffer (75mM KCl, 50mM Tris pH8.3, 3mM MgCl2, 10mM DTT) with 4% glycerol and 20μg/ml BSA prior to immunoprecipitation; the RNase H treatment was performed for 2 hours at 37°C.

For comparing POLR2A and S9.6 signals on the *ACTB* gene, wild type or control signals were normalized to 1, and the R1810A mutant, knock-down or knock-out samples were adjusted such that the ratio for the intron 3 (1671) position was set to 1. Similarly, for the *GAPDH* gene, the ratio for the intron 5 (2436) position was set to 1. ChIP data for SMN, FUS, and TDP-43 were expressed as percent input or as ratio to the ChIP data for POLR2A. Error bars represent biological replicates, except where indicated otherwise.

### Primer information

Primers used in ChIP are listed here (15,37). For the *ACTB* gene:

-72.fw CCGAAAGTTGCCTTTTATGGC, -72.rev CAAAGGCGAGGCTCTGTGC;

332.fw CGGGGTCTTTGTCTGAGC, 332.rev CAGTTAGCGCCCAAAGGAC;

1671.fw TAACACTGGCTCGTGTGACAA, 1671.rev AAGTGCAAAGAACACGGCTAA;

2911.fw TGCGCAGAAAACAAGATGAG, 2911.rev GTCACCTTCACCGTTCCAGT;

3560.fw TTACCCAGAGTGCAGGTGTG, 3560.rev CCCCAATAAGCAGGAACAGA;

3752.fw GGGACTATTTGGGGGTGTCT, 3752.rev TCCCATAGGTGAAGGCAAAG;

4657.fw TGGGCCACTTAATCATTCAAC, 4657.rev CCTCACTTCCAGACTGACAGC;

5590.fw CAGTGGTGTGGTGTGATCTTG, 5590.rev GGCAAAACCCTGTATCTGTGA.

For the *GAPDH* gene: 55.fw CTCCTGTTCGACAGTCAGC, 55.rev TTCAGGCCGTCCCTAGC;

1407.fw CACCCTGGTCTGAGGTTAAATATAG, 1407.rev GTGGGAGCACAGGTAAGT;

2436.fw ATAGGCGAGATCCCTCCAA, 2436.rev TGAAGACGCCAGTGGAC;

3882.fw CCCTGTGCTCAACCAGT, 3882.rev CTCACCTTGACACAAGCC;

4511.fw AGATGTGTCAGGGTGACTTAT, 4511.rev TAGGTCCCAGCTACACGC;

5196.fw GTCTCAGTGTATGACAGACACG, 5196.rev TGTATGTGCGCTCAGGG.

### Chromatin immunoprecipitation and Sequencing analysis (ChIPseq)

Chromatin immunoprecipitation was performed as before (38). In brief, 10^7^-10^8^ cells were cross-linked for 10 min in 1% formaldehyde. Lysates were sonicated to a DNA fragment length range of 200-300 bp using a Bioruptor (Diagenode). RNAPII was immunoprecipitated with 2ug of antibodies and Dynabeads Protein G (Invitrogen). Subsequently, crosslinks were reversed at 65°C over night and bound DNA fragments were purified (EZ-10 Spin Column PCR Product Purification kit, Bio Basic). Sequencing libraries were constructed using the TruSeq ChIP Sample Prep Kit (Illumina) according the manufacturer’s instructions. Libraries were sequenced (single end reads) on the Illumina HiSeq 2500 to a minimum depth of 20 million reads, obtaining at least 12-20 million unique reads per sample.

ChIPseq analysis was performed as before for the display of meta-gene plots (9). Reads in FASTQ format were mapped to the human genome (hg19) using Bowtie 2 (39) with options −5 1 −3 2 –local duplicate reads removed, and reads were extended to 300 bp. The number of fragments overlapping each genomic bp were calculated and were normalized by million mappable reads in the ChIP-seq library. Only the top 5 or 10% of genes by mRNA expression levels were considered.

For the calculating of promoter stalling and termination stopping ratios, signal density was calculated using the SPP (40) R package (bandwidth 150bp, step 50bp). Background from input was first scaled based on sequencing depth (total number of unique reads) and then subtracted from sample signal. Signal density in different experimental groups was quantile normalized to allow direct of comparison between mutants and wild-type controls. Promoter region was defined as −30bp to + 300bp around the transcription start site. The rest (from +301 to the end of annotated gene) was defined as the gene body. Termination region was defined as the 2000bp segment downstream from the polyA cleavage site. Signals in each of these regions were summed up and divided by the length of the region to get a normalized value. Promoter stalling ratio was defined as the ratio of promoter region signal over gene body signal, and termination stopping ratio was defined as the ratio of termination region signal over gene body signal. Only genes with signal in gene body greater than 1 were included in analysis. Quantiles of promoter stalling and termination stopping ratios were computed for KO mutants and control separately and plotted against increasing level of ratios. Statistically significant differences between the quantiles were determined using the Kolmogorov Smirnov test; level of significance is set at p < 0.05. Only top 10% genes by mRNA expression levels were considered; statistical analysis and data plotting were performed using R.

### Antibodies, constructs and reagents

Anti-CTD R1810me2s antibody was raised in rabbits using a KLH-conjugated CTD peptide from POLR2A (amino acids 1806-1813) that carried an R1810me2s modification. KLH conjugation was performed using an N-terminal cysteine residue (Cedarlane) (9). R1810me2s-specific antibodies were enriched by flowing the serum through a column containing an R1810me0 peptide conjugated to SulfoLink coupling resin (Thermo Scientific, 20401).

Flp-in TREx GFP-HB fusion construct that contains the R-loop binding domain of RNase H was provided by Dr. Andres Aguilera (41). The ORFs for TDP-43 came from the Plasmid collection at Harvard. GFP constructs were generated via Gateway cloning into pDEST pcDNA5/FRT/TO-eGFP. α-amanitin resistant wild type and R1810A mutant POLR2A constructs were kindly provided by Dr. Dirk Eick (8); RNAPII R1810me2a antibody was kindly provided by Dr. Danny Reinberg (8). We obtained the POLR2A pSer2 and pSer5 antibodies from the Eick laboratory (S2P: 3E10; S5P: 3E8). 8WG16 antibody against unphosphorylated CTD repeats of POLR2A was prepared in the lab.

Commercial antibodies were as follows: HA (Sigma, mAb H9658); PRMT5 (Upstate, pAb C7-405, Santa Cruz, mAb sc-22132); SMN (Santa Cruz, pAb H-195); SETX for ChIP and IP (Novus Biologicals, pAb NB100-57543) and for western blots (Bethyl Lab, pAb A301-104A); XRN2 (Santa Cruz, pAb sc-99237); FUS (Santa Cruz, mAb sc-47711); TDP-43 (Bethyl lab, pAb A303-233A); POLR2A N20 (Santa Cruz, pAb sc-899); POLR2A 4H8 (Abcam, mAb ab5408); gammaH2Ax (Millipore, 05-636); H2Ax (Millipore, 07-627); Tubulin (Sigma, mAb T8328); GFP (Abcam, pAb 290); IgG negative controls for ChIP and IP (Millipore, pAb 12-370). α-amanitin was purchased from Sigma (23109-05-9).

## RESULTS

### 1. Recruitment of TDP-43 to RNAPII is mediated by SMN and CTD R1810

We investigated the roles of FUS and TDP-43 with respect to the RNAPII CTD R1810me2s-SMN pathway, as both are arginine-dimethylated proteins (20,42) that interact with each other and with SMN (43,44) and are sometimes mutated in the same neurodegenerative disease, ALS. We first used immunostaining to show that FUS and TDP-43 are localized in the nuclei of HEK293 cells, as are the phosphorylated forms of POLR2A and the RNAPII termination factors XRN2 and SETX (Supplementary Figure S1). Next, from multiple co-IP experiments, we observed that FUS and TDP-43 interact directly or indirectly with SMN and POLR2A, as well as termination factors such as SETX and XRN2 (Figure 1A), as summarized in Figure 1B. Many of these interactions occur independently of RNAPII, as they persist after a 3-day α-amanitin treatment of HEK293 cells that eliminated the bulk of the RNAPII (Supplementary Figure S2A). It is likely that FUS, TDP-43, XRN2, and SETX are PRMT5 substrates, because they all interact with PRMT5 (Figure 1A) and are known to contain dimethylated arginine (Supplementary Table 1).

**Figure 1.**
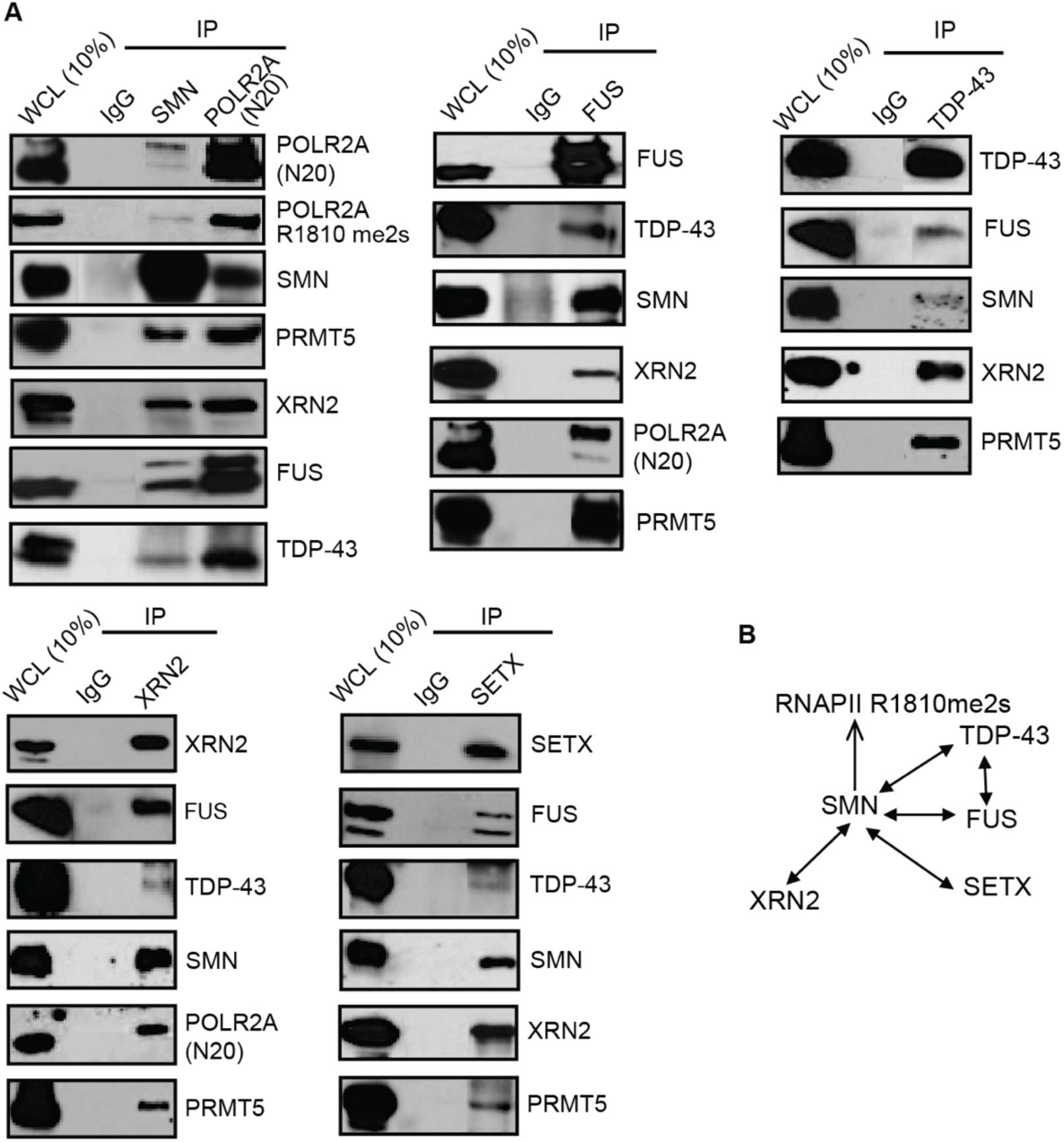
Interaction network among proteins involved in transcription termination. **A**. Immunoprecipitation (IP) with the indicated antibodies from HEK293 whole cell lysate (WCL), followed by western blotting with the indicated antibodies. IP with IgG was used as a negative control. The blots indicate that POLR2A, SMN, FUS, TDP-43, PRMT5 and the termination factors SETX and XRN2 interact directly or indirectly with each other (9). **B**. A summary of the interactions detected by co-IP experiments. All of these interactions, except the SMN-XRN2 and SMN-TDP-43 interactions, have been shown by various published experiments to be direct (9,14,35).

Chromatin immunoprecipitation (ChIP) was performed for FUS and TDP-43, and the data for the *ACTB* gene (Figure 2A) are represented in Figure 2B-C as percent input or as ratio with respect to RNAPII. These experiments showed that FUS and TDP-43 associate with the *ACTB* gene throughout its length from the promoter to the termination regions. We have previously shown that the RNAPII CTD R1810me2s modification recruits SMN to enhance the interaction of SETX with RNAPII; here we show that the association of TDP-43 and, to a lesser extent, FUS with RNAPII is similarly enhanced by the presence of the RNAPII CTD R1810 residue *in vivo*. First, we observed that the interactions of TDP-43 and FUS with HA-POLR2A were reduced when R1810 was mutated to alanine on the POLR2A CTD (Figure 2D, Supplementary Figure S2B). These experiments were performed in Raji cells after 3 days of α-amanitin treatment that eliminated the bulk of the endogenous RNAPII (9). Interactions were detected by immunoprecipitating α-amanitin-resistant, HA-tagged, wild type or R1810A mutant POLR2A using anti-HA antibodies. Additionally, by immunoprecipitating RNAPII, we found that the interaction between POLR2A and TDP-43 was consistently reduced when SMN or PRMT5 was knocked down or when SMN was knocked out using CRISPR/Cas9-mediated mutagenesis in HEK293 cells (9) (Figure 3A-B). Since FUS is able to associate directly with the RNAPII CTD (45,46) *in vivo* and *in vitro*, the interaction of FUS with RNAPII was also observed to be independent of SMN in some cases (Figure 3B).

**Figure 2.**
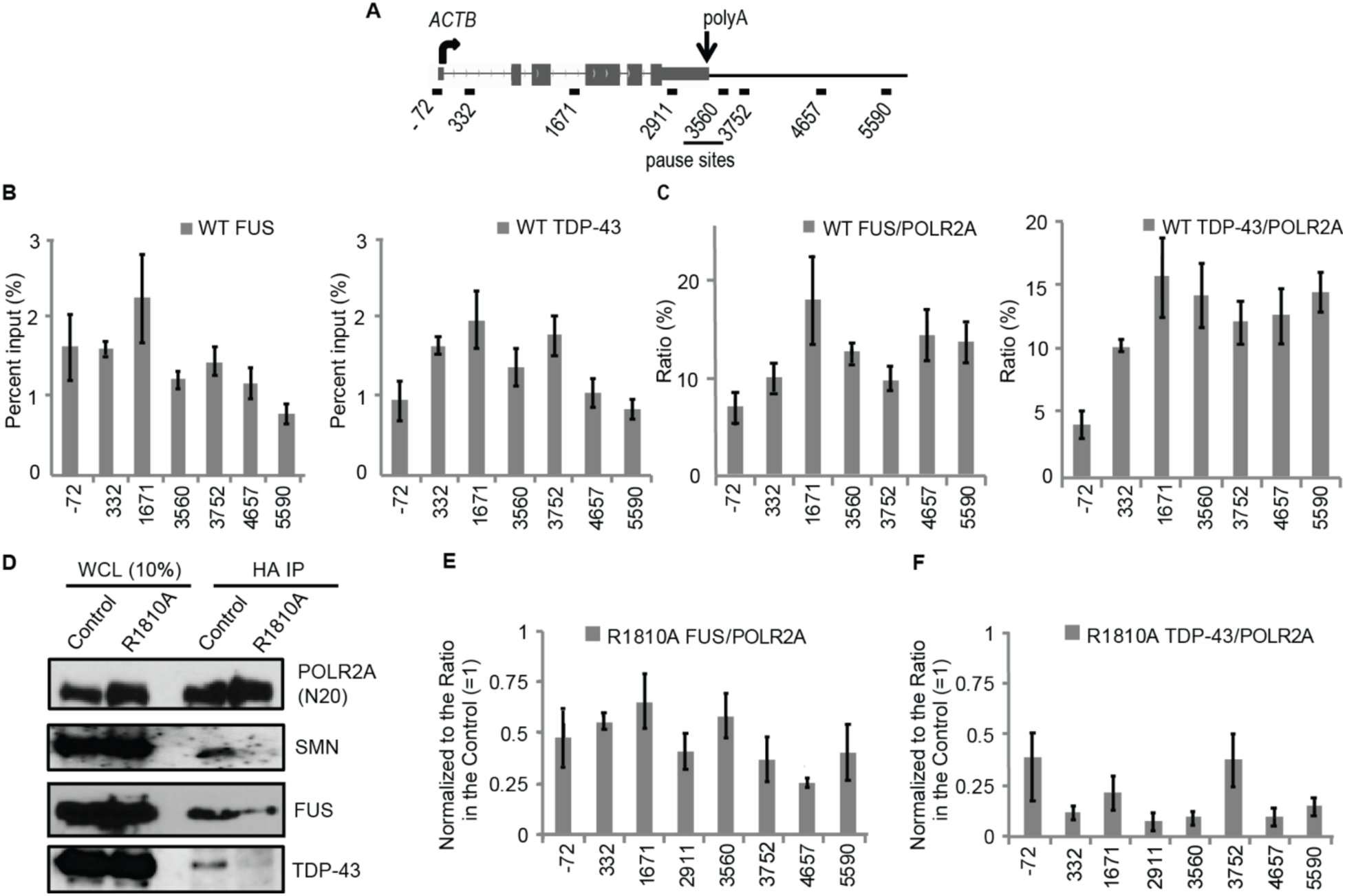
R1810 on the RNAPII CTD enhances the recruitment of SMN, FUS, and TDP-43. **A**. Primer positions along the *ACTB* gene; the termination pause sites are underlined. **B, C**. Quantification of ChIP data in HEK293 cells, expressed as percent input, or as ratio with respect to RNAPII, using the indicated primer positions for qPCR along the *ACTB* gene. Error bars denote biological replicate s.d. (n=3). **D**. IP with the indicated antibodies from WCL from Raji cells stably expressing α-amanitin-resistant HA-tagged wild-type or R1810A mutant POLR2A after α-amanitin treatment that abolishes the endogenous POLR2A. Anti-HA was used to precipitate HA-tagged wild-type (Control) or R1810A mutant POLR2A, followed by western blotting with the indicated antibodies. **E, F**. Quantification of ChIP data in Raji cells as FUS/POLR2A (**E**) or TDP-43/POLR2A (**F**) ratio to show the relative effects of mutating R1810 to alanine, with the signals in wild type R1810 controls normalized to 1. Error bars denote biological replicate s.d. (n=3).

**Figure 3.**
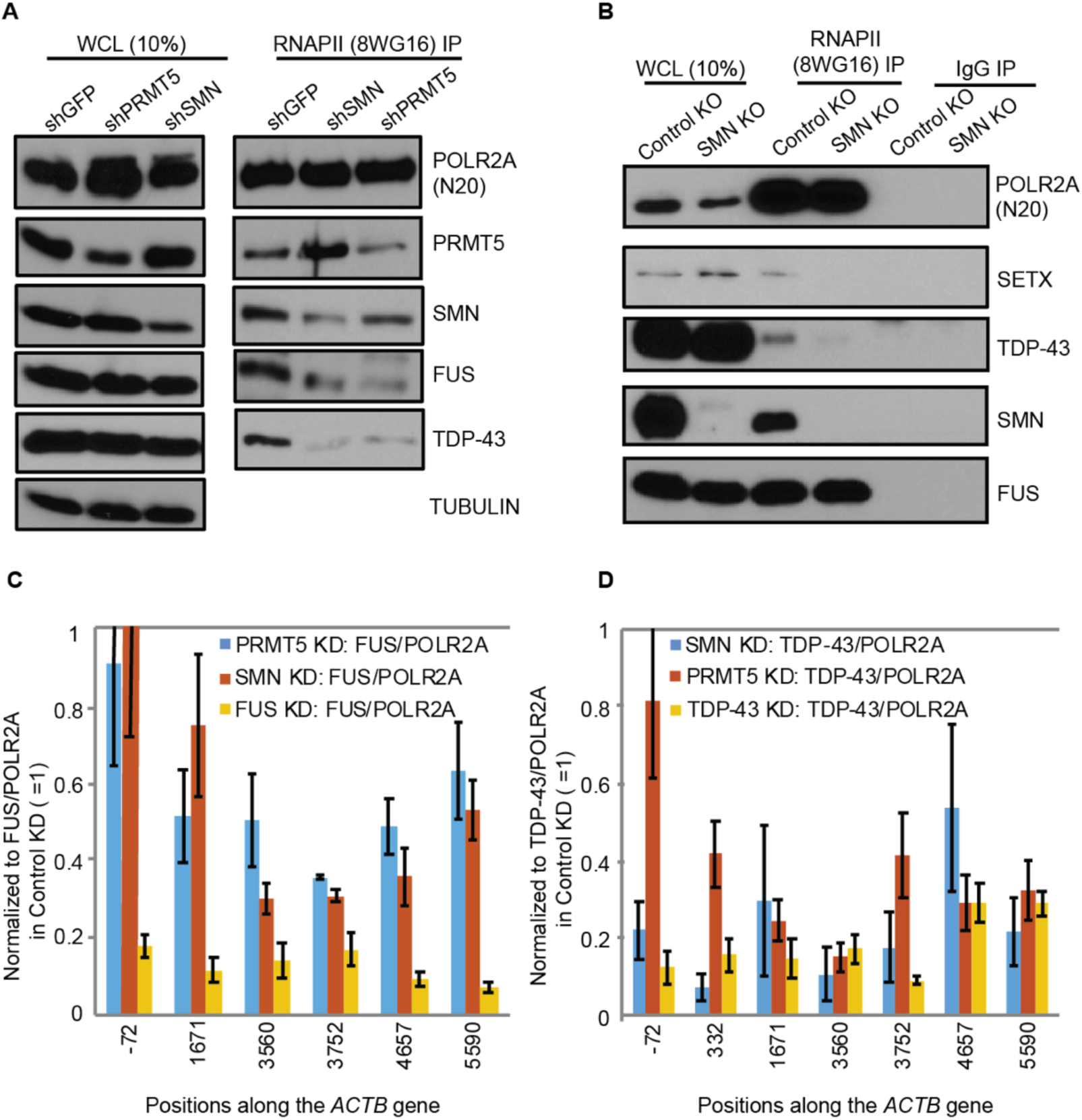
The loss of PRMT5 or SMN disrupts the interactions among RNAPII, SMN, FUS, and TDP-43. **A**. IP with 8WG16 antibodies for POLR2A from HEK293 WCL upon stably knocking down PRMT5 or SMN, with the knock-down of GFP as negative control. Western blots were performed with the indicated antibodies. The knock-down of PRMT5 or SMN caused a reduction of FUS and TDP-43 interaction with RNAPII. **B**. IP with the indicated antibodies from HEK293 WCL, upon knocking out of SMN with the CRISPR/Cas9 system or using scrambled guide RNA as negative control. Western blots were performed with the indicated antibodies to show that the SMN knock-out leads to a loss of interaction of TDP-43 and SETX with POLR2A. IP with IgG as negative control. **C, D**. Quantifications of ChIP in HEK293 cells as FUS/POLR2A (**C**) or TDP-43/POLR2A (**D**) ratio to show the relative effects of knocking down PRMT5 or SMN. Error bars denote biological replicate s.e.m. (n=3).

To further characterize the roles of CTD R1810me2s in recruiting TDP-43 or FUS to RNAPII, ChIP experiments were performed. ChIP involving α-amanitin-resistant, wild type or R1810A POLR2A mutant cell lines was done after 3 days of treatment with α-amanitin (2ug/ml) to transiently deplete the endogenous RNAPII. The FUS/POLR2A and TDP-43/POLR2A ChIP signals along the length of the *ACTB* gene were reduced in the RNAPII R1810A mutant, especially for TDP-43 (Figure 2E-F). Upon the knock-down of PRMT5 or SMN, a reduced association of TDP-43 and FUS with transcribing POLR2A was also observed in ChIP experiments (Figure 3C-D). The specificities of the FUS and TDP-43 antibodies in ChIP assays were verified upon the knock-down of either protein for ChIP (Figure 3C-D). Collectively, these observations suggest that the association of TDP-43 and, to a lesser extent, FUS with RNAPII is enhanced by SMN and RNAPII CTD R1810me2s.

### 2. Endogenous RNAPII R1810A mutant is defective in RNAPII termination

In our previous studies (9) and in Figure 2, we used α-amanitin to eliminate the endogenous RNAPII to assay the transient effect of the R1810A mutation on the α-amanitin-resistant HA-tagged POLR2A. To examine the stable effect of an R1810A mutation, CRISPR/Cas9-mediated knock-in of the mutation in the endogenous POLR2A gene was generated in HEK293 cells. We checked by DNA sequencing that R1810 on both alleles of the endogenous POLR2A were mutated to alanine (not shown). When we precipitated wild type and R1810A mutant RNAPII using antibodies against the hypophosphorylated IIA (8WG16) form of the CTD (Figure 4A), the R1810A mutant RNAPII showed a loss of signal for both R1810me2a and R1810me2s modifications, as detected by their specific antibodies (8,9). We previously found that CTD phosphorylation interfered with detection of R1810me2s and, consistent with that, R1810me2s was better detected only when we precipitated hypophosphorylated RNAPII (not shown). We then performed RNAPII ChIP and found that the R1810A mutant accumulated more in the termination regions of the *ACTB* and the *GAPDH* genes, in agreement with our observations with the α-amanitin-resistant HA-POLR2A in Raji cells (Figure 4B, Supplementary Figure S3A) (9).

**Figure 4.**
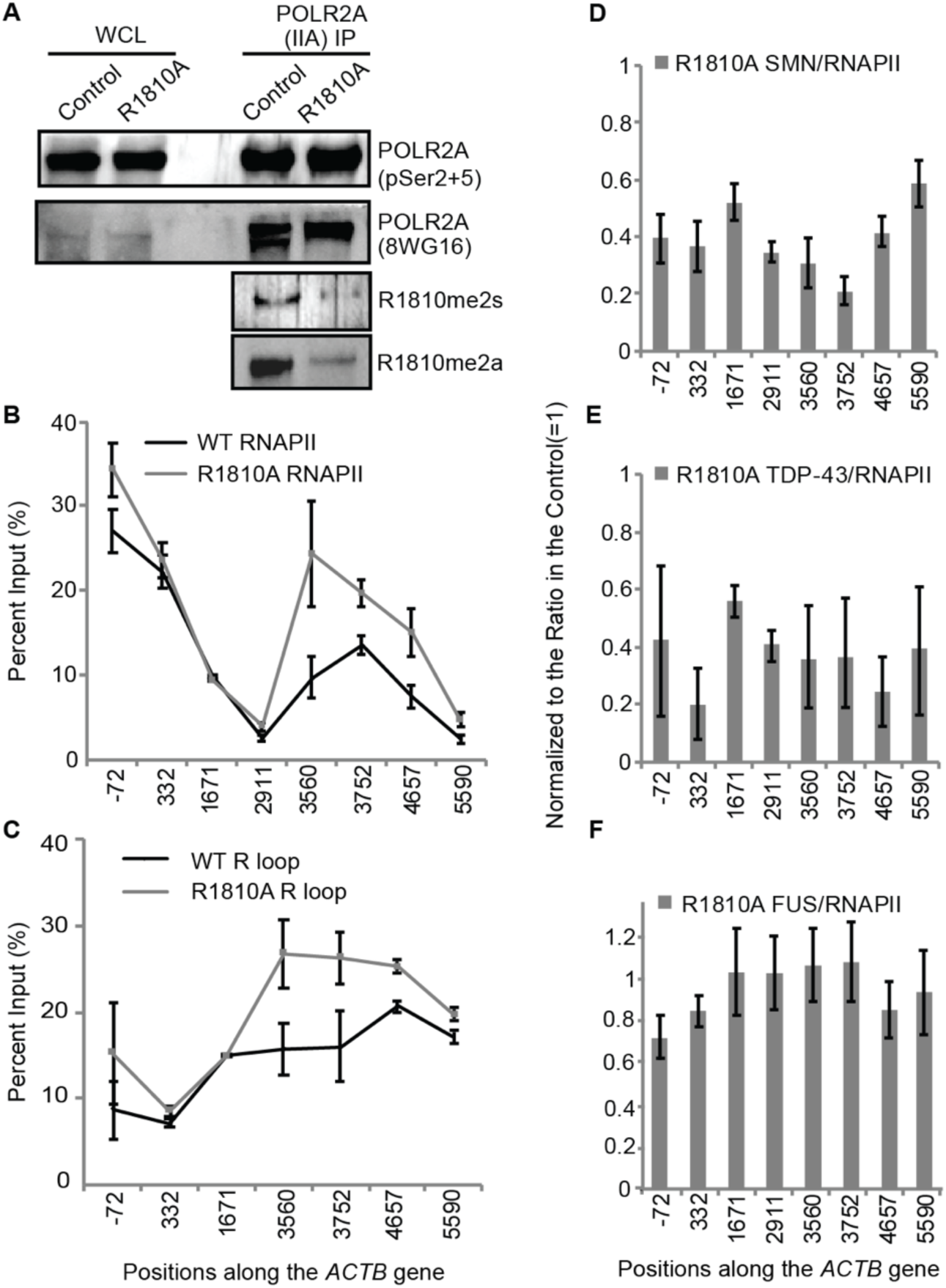
Endogenous RNAPII CTD R1810A mutant also causes an RNAPII termination defect and R-loop accumulation, as well as reduced recruitment of SMN and TDP-43. **A**. Immunoprecipitation (IP) with the indicated antibodies from HEK293 whole cell lysate (WCL), followed by western blotting with the indicated antibodies. The blots indicate that the R1810A mutation on RNAPII causes the loss of both R1810me2a and R1810me2s signals as detected by specific antibodies. **B**. Quantification of RNAPII ChIP for WT or the R1810A mutant POLR2A with antibodies (8WG16, N20) in HEK293 cells, using the indicated primer positions for qPCR along the *ACTB* gene. Error bars denote biological replicates as s.e.m. (n=5). **C**. Quantification of R-loops with the S9.6 antibody in HEK293 cells for the WT or the R1810A mutant POLR2A, using the indicated primer positions for qPCR along the *ACTB* gene. Error bars denote biological replicates as s.e.m. (n=4). **D, E, F**. Quantification of ChIP data in HEK293 cells as SMN/POLR2A (**D**), TDP-43/POLR2A (**E**), or FUS/POLR2A (**F**) ratio to show the relative effects of mutating R1810 to alanine, with the signals in the wild type R1810 control normalized to 1. Error bars denote biological replicate s.d. (n=3).

We and others have shown that the resolution of R-loops formed by elongating RNAPII in its termination regions is enhanced by RNAPII CTD R1810me2s, SMN, and SETX (9,15). The monoclonal antibody S9.6 detects R-loops in DNA immunoprecipitation experiments, and the signals were shown to be specific upon RNase H treatment (Supplementary Figure S4A) (9). Here, we show that the R1810A mutation on the endogenous RNAPII also increased R-loop accumulation in the termination regions of the *ACTB* (Figure 4C) and *GAPDH* (Supplementary Figure S4B) genes. The regions where RNAPII stalled correlated well with R-loop accumulation in the R1810A mutant. We further carried out ChIP of SMN, TDP-43, and FUS along the length of the *ACTB* gene, and observed that the ChIP signals for SMN and TDP-43 with respect to RNAPII were reduced in the R1810A mutant as compared to the wild type (normalized to 1) (Figure 4D-E), similar to the observation made with the α-amanitin resistant HA-tagged POLR2A (Figure 2F) (9). However, FUS ChIP signals showed less or no reduction, likely due to alternative or compensatory mechanisms for its interaction with the RNAPII CTD (Figure 4F).

### 3. FUS and TDP-43 are important for transcription termination by RNAPII

As SETX, SMN and the RNAPII CTD R1810me2s are important for R-loop resolution and termination by RNAPII (9,15), we investigated if similar effects can be observed upon the loss of FUS or TDP-43. Upon stable shRNA-mediated knock-down or CRISPR/Cas9-mediated knock-out of FUS or TDP-43 in HEK293 cells (Supplementary Figure S3B), RNAPII indeed showed accumulation in the termination regions of the *ACTB* (Figure 5A, Supplementary Figure S5) and *GAPDH* (Supplementary Figure S3C-D) genes. FUS and TDP-43 contain RNA recognition motifs (RRMs) that could potentially bind to nascent RNA and prevent its hybridization with DNA for the formation of R-loops. To test this idea, we used HEK293 Flp-In T-REx cell lines to express a GFP-tagged wild-type TDP-43 or a truncated mutant that lacks its two RNA binding domains (RRMs) (Supplementary Figure S6A-B). With RNAPII ChIP signals normalized in gene bodies to a control that over-expresses wild type TDP-43, the over-expression of the truncation mutant led to increased accumulation of RNAPII in the termination regions of the *ACTB* (Figure 5B) and *GAPDH* (Supplementary Figure S6C) genes. This suggested that the RNA-binding ability of TDP-43 is important for its role in RNAPII termination and R-loop resolution, perhaps through sequestering the nascent RNA from hybridizing to single-stranded DNA behind the transcription bubble.

**Figure 5.**
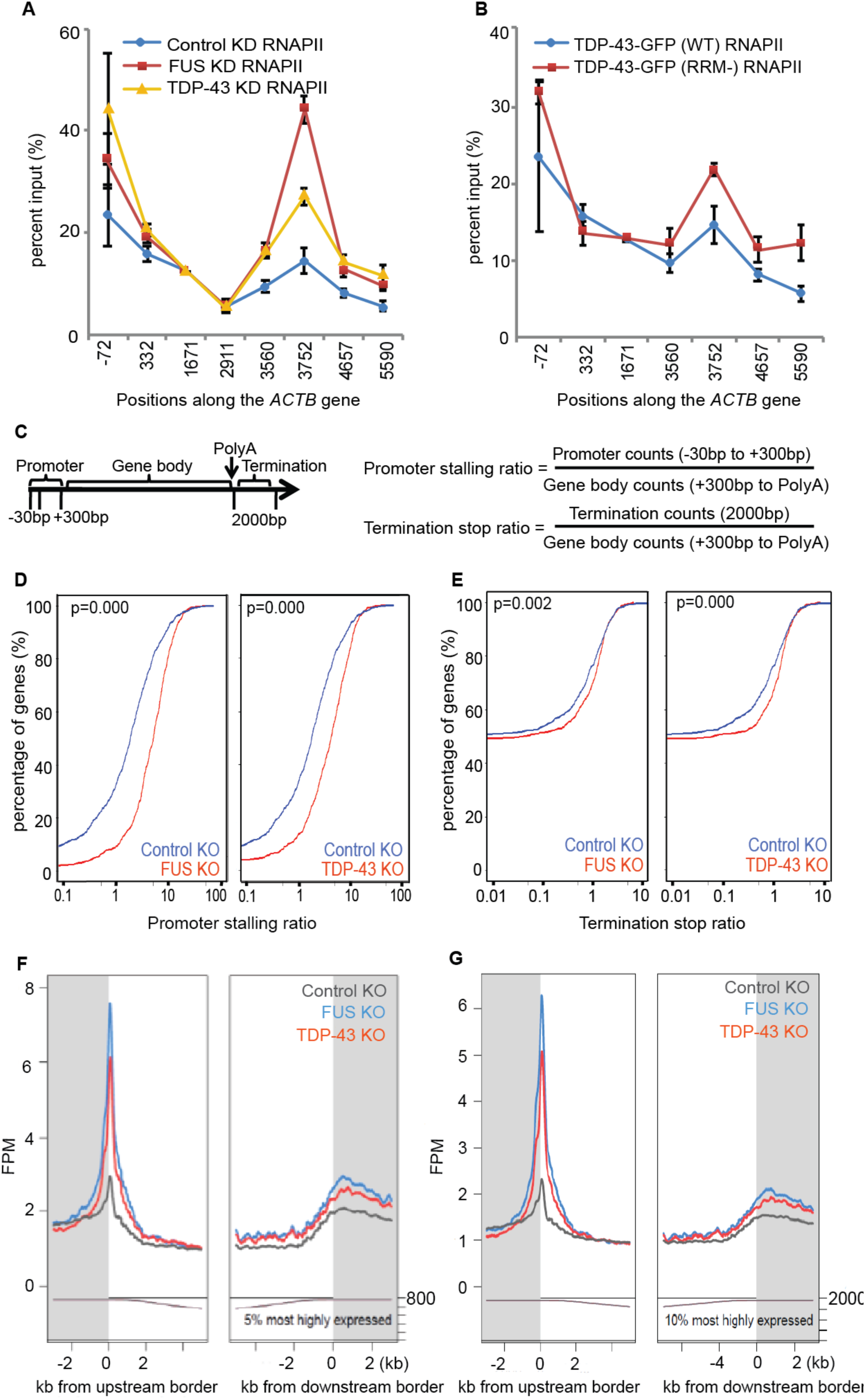
FUS and TDP-43 regulate transcription termination by RNAPII. **A**. Quantification of RNAPII ChIP using POLR2A antibody (4H8, N20) in HEK293 cells, using the indicated primer positions for qPCR along the *ACTB* gene after stably knocking down FUS or TDP-43, with GFP knock-down as a negative control. Error bars denote biological replicates as s.e.m. (n = 3-5). **B**. Quantification of RNAPII ChIP using the POLR2A antibody (N20) in HEK293 cells, with cells overexpressing wild type GFP-TDP-43 or the truncation mutant lacking RNA binding domains GFP-TDP-43 (RRM-), using the indicated primer positions for qPCR along the *ACTB* gene. Error bars denote biological replicates as s.e.m. (n = 3). **C**. Schematics for the calculation of the stalling ratio at the promoter region and stop ratio at the termination region. A rightward shift of the curve in the plots indicates increased RNAPII accumulation in the corresponding regions. **D, E**. RNAPII ChIP-seq (with the N20 antibody in HEK293 cells). Compared to the scrambled KO, FUS or TDP-43 KO led to increased RNAPII accumulation in the promoter (**D**) and termination (**E**) regions of many genes. Top 10% genes by expression were analyzed. Statistically significant differences between the quantiles were determined using the Kolmogorov Smirnov test; level of significance is set at p < 0.05. **F, G**. A second way of displaying ChIP-seq analysis in which the reads are normalized to the gene body. The knock-out of FUS or TDP-43 led to increased RNAPII accumulation in the promoter and termination regions of many genes. Top 800 (**F**) or 2000 (**G**) genes by expression with reliable ChIP-seq signals were analyzed; the number of genes analyzed is displayed on the bottom right of the graph.

We then performed RNAPII ChIP-seq to determine whether, as with the effects of loss of SMN or the presence of an R1810A mutation in the RNAPII CTD, the loss of FUS or TDP-43 had general effects on transcription termination. Using HEK293 cells with a CRISPR/Cas9 knock-out of FUS or TDP-43, or with a scrambled guide RNA as control, RNAPII ChIP-seq (with the N20 antibody) was performed with the generation of 12-20 million unique reads in each case. For the analysis, we used genes with reliable ChIP-seq signals (top 5 or 10% by expression; approximately 800 or 2000 genes, respectively), as sequencing depth and background interference became issues when more genes were included. Quantile plots were generated for the promoter-stalling ratio and the terminator stop ratio, calculated as promoter region reads normalized to gene body reads, or terminator region reads normalized to gene body reads, respectively (Figure 5C-E). A rightward shift of the plot indicates increased RNAPII accumulation in the corresponding regions (Figure 5C-E). The lack of FUS or TDP-43 caused substantial accumulation of RNAPII in promoter regions, in agreement with the literature for the loss of FUS (Figure 5D, F, G) (27,45). Supporting the roles of FUS and TDP-43 in the regulation of RNAPII termination, we observed genome-wide accumulation of RNAPII in the termination regions when either protein was knocked out (Figure 5E-G; Supplementary Figure S7).

### 4. FUS and TDP-43 prevent the accumulation of R-loops and DNA damage at termination sites

Using the S9.6 antibody, which binds RNA/DNA hybrids, for R-loop detection (9,15), we found that, similarly to the effect of an RNAPII R1810A mutation or the loss of SMN or SETX, the knock-down of FUS or TDP-43 also led to R-loop accumulation in the termination regions of the *ACTB* gene (Figure 6A) where RNAPII stalls. To further address the validity of R-loop detection, we employed a second way of R-loop detection using a stably expressed GFP fusion protein that includes the RNase H1 R-loop-binding domain (GFP-HB) (41). Indeed, increased accumulation of R-loops was detected by anti-GFP ChIP for GFP-HB in the termination regions of the *ACTB* (Figure 6B) and *GAPDH* (Supplementary Figure S4C) genes when FUS or TDP-43 was knocked out. These experiments indicated that RNAPII CTD R1810me2s, SMN, SETX, FUS, and TDP-43 all regulate R-loop resolution and transcription termination by RNAPII.

**Figure 6.**
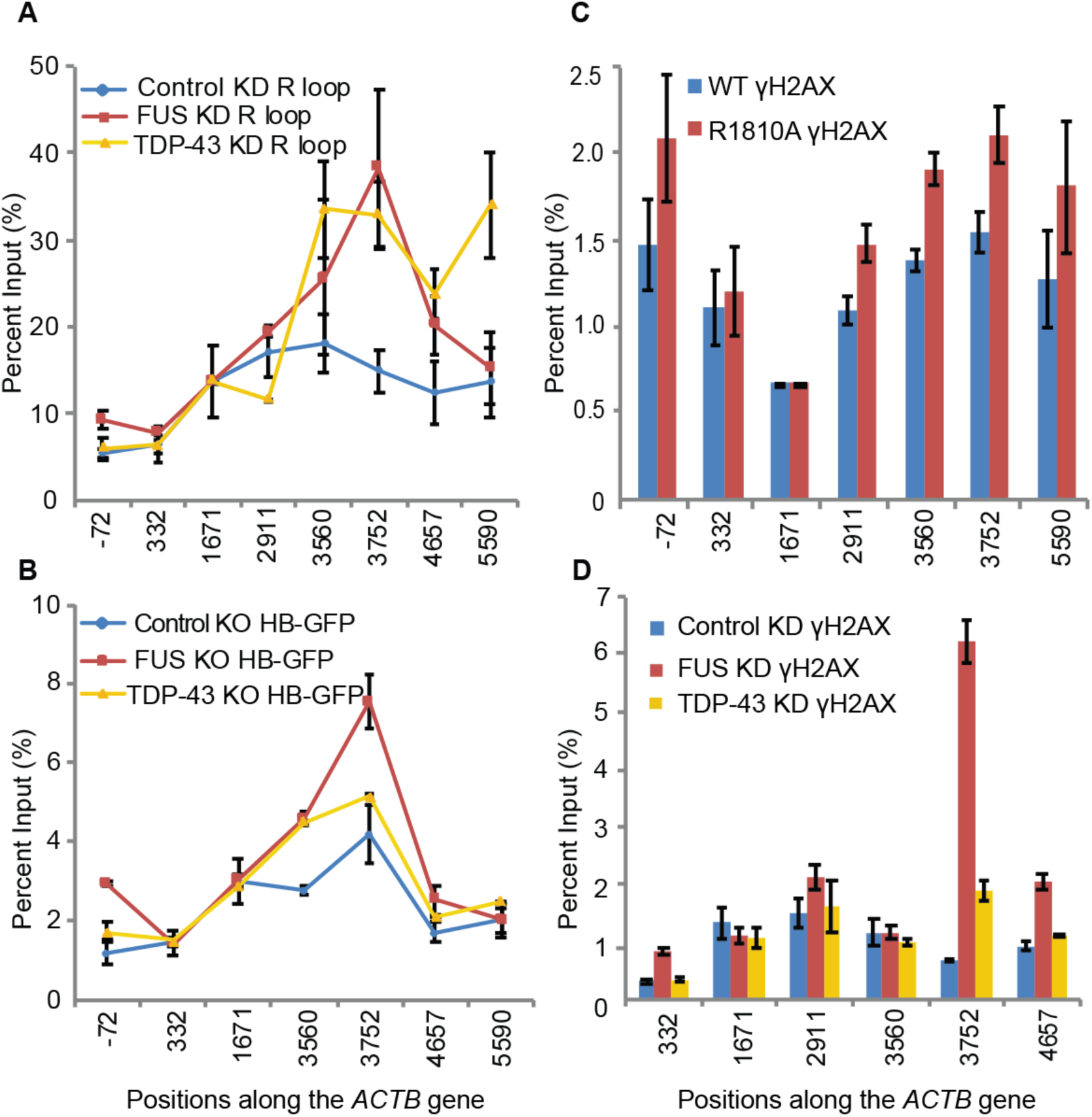
FUS and TDP-43 prevent accumulation of R-loops and DNA damage at RNAPII transcription terminators. **A**. Quantification of R-loops with the S9.6 antibody in HEK293 cells with or without knock-down of FUS or TDP-43, using the indicated primer positions for qPCR along the *ACTB* gene. Error bars denote biological replicates as s.e.m. (n=4). **B**. ChIP quantification of R-loop with the GFP-HB construct, with the indicated primer positions for qPCR along the *ACTB* gene, after knocking out FUS or TDP-43 by CRISPR/Cas9 mutagenesis, using scrambled guide RNA as a negative control. Error bars denote biological replicates as s.e.m. (n=3). **C, D**. ChIP quantification of γH2AX as percent input in HEK293 cells, along the length of the *ACTB* gene, between WT and the endogenous R1810A mutant POLR2A (**C**), or after knocking down FUS or TDP-43 (**D**). Error bars denote biological replicates as s.e.m. (n=4).

Because R-loop accumulation during transcription is associated with DNA damage and genome instability (9,47-54), we also examined the level of DNA damage at terminators upon loss of FUS or TDP-43 by using γH2AX ChIP as a proxy for the accumulation of DNA damage (9,55) (Figure 6C-D; Supplementary Figure S8). We indeed observed an accumulation of γH2AX and increased γH2AX/H2AX ratio in the termination regions of *ACTB*, where RNAPII and R-loops also accumulated upon the knock-down of FUS or TDP-43, or upon mutating R1810 to alanine in the endogenous POLR2A. Therefore, our study brings further support to the idea that RNAPII and R-loop accumulation in termination regions could lead to genome instability that contributes to neurodegenerative pathology (9).

## DISCUSSION

SMN’s ability in self-associate (12,56) is central to its function as an adaptor for the assembly of the spliceosome and cytoplasmic ribonucleoprotein complexes, defects in which are known to cause SMA neuromuscular pathology (11). As we found that SMN interacts with multiple termination factors that contain dimethylated arginine (e.g. FUS, TDP-43, XRN2), an additional pathway that SMN may coordinate is the assembly of an R-loop resolution complex to regulate RNAPII termination (Figure 7A-B). Since SMN, FUS, and TDP-43 form protein-protein interactions, it has been argued that the ALS and SMA diseases are molecularly related diseases (43,44). Among the ∼150 human proteins that have been shown by mass spectrometry to contain dimethylated arginine, more than 50 of them are involved in RNAPII transcription and termination (16-20). Besides POLR2A (9), various well-known transcriptional elongation and termination factors such as FUS, TDP-43, EWSR1, TAF15, CPSF1, CPSF5, CPSF6, SPT5, CTDP1, PABP1, and PABP2 all contain arginine dimethylations (Supplementary Table 1) (17).

**Figure 7.**
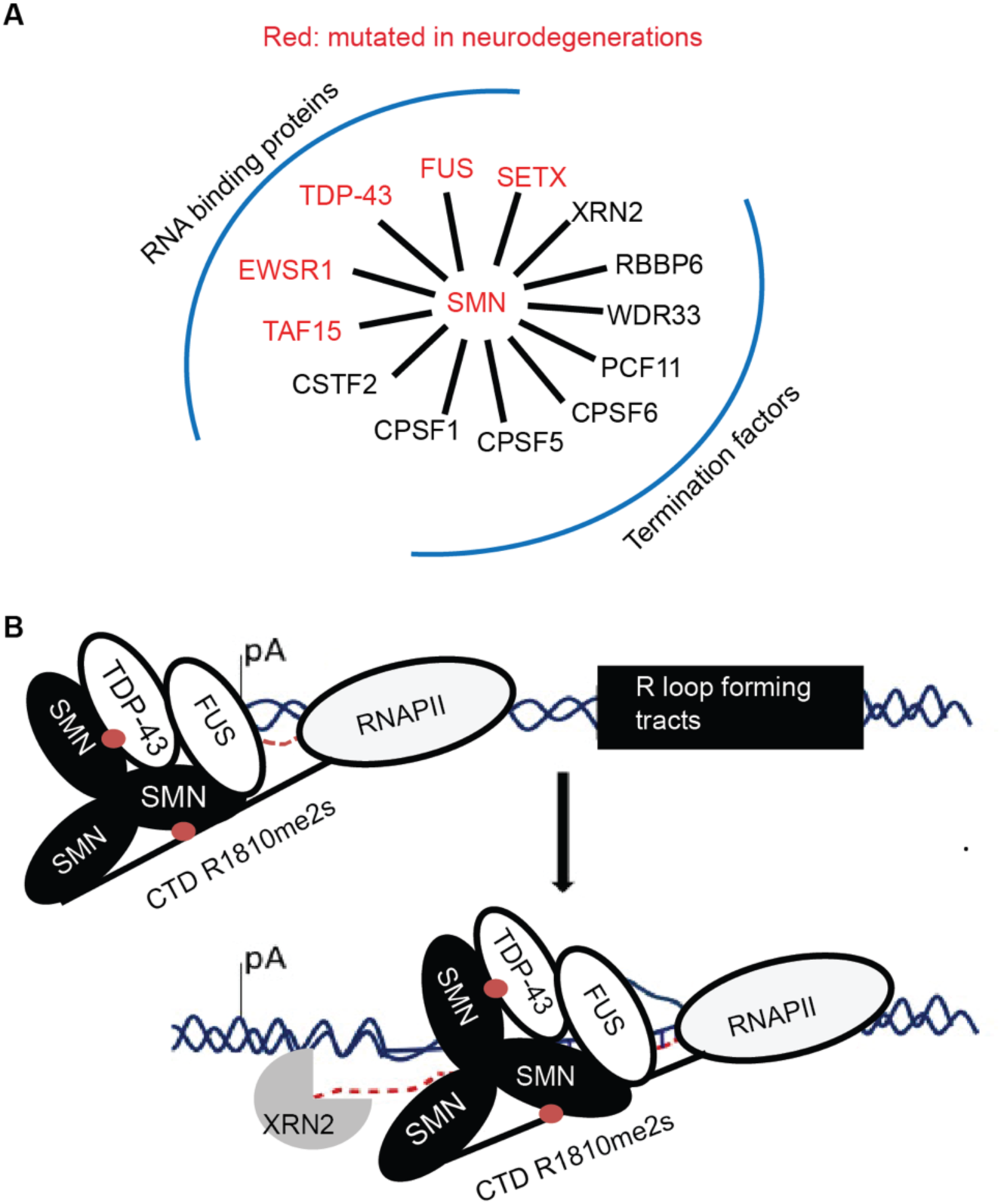
Model. SMN as a nucleator of arginine dimethylated proteins involved in transcription termination and neurodegeneration (**A**), through the recruitment of FUS or TDP-43 to prevent R-loop formation and DNA damage at RNAPII termination sites (**B**).

It is known that the presence of GC- or G-rich tracts at a locus (15,57) can induce R-loop accumulation more readily, and defects in R-loop elimination are linked to genome instability (35,47,55). G-rich DNA tracts can frequently be found in promoters and termination regions, as well as regions with trinucleotide repeat expansions that are responsible for various neuronal diseases (e.g. ALS/FTD, Fragile X syndrome, and Friedreich’s Ataxia) (58-61). Such tracts are known to induce DNA damage and mutations, as they are often stabilized upon the knock-down of certain factors involved in the transcription-coupled nucleotide excision repair (48), homologous recombination (62), and the Fanconi Anemia pathway (53,63). It is also known that multiple surveillance mechanisms in eukaryotes exist to eliminate the buildup of R-loops and the consequent DNA hyper-recombination (64).

The ability of FUS and TDP-43 to bind RNA, thereby potentially preventing the formation of R-loops, may explain why we observed increased RNAPII accumulation in termination regions when either FUS or TDP-43 is knocked out (Figure 7). FUS is known to be associated with the RNAPII CTD and to regulate the phospho-Ser2 level of the CTD, and it is also known to promote RNAPII termination and alter the site of cleavage and polyadenylation of nascent RNAs (27,45) with a preference for binding the G-rich sequence GGUG (65). TDP-43, on the other hand, binds to GU-rich sequences and can compete with termination factors for UG-rich sequences downstream of the polyadenylation signal (66,67). Furthermore, both FUS and TDP-43 are known to suppress the buildup of transcription-associated DNA damage (52,68-72). Given that ALS or FTD develops after decades of life, we hypothesize that the accumulation of DNA damage due to the buildup of R-loops at transcription terminators in post-mitotic neurons contributes to neurodegenerative pathology.

## DATA AVAILABILITY

Data for ChIP-seq analyses have been deposited in GEO with the accession code GSE134332.

## SUPPLEMENTARY DATA

Supplementary Data are available at NAR online.

## ACKNOWLEDGEMENT

We thank D. Reinberg for RNAPII R1810me2a antibodies; D. Eick for wild-type and RNAPII (R1810A) constructs; and S. Leppla for purified S9.6 antibodies. We also thank D. Torti and D. Leung for Illumina library preparation and sequencing, J. Li for purified 8WG16 antibodies, and D. Durocher for constructive criticism and advice during the course of this work.

## Author Contributions

J.F.G. supervised the project. D.Y.Z. performed the experiments. J.F.G. and D.Y.Z. wrote the manuscript. B.J.B and Z.N commented on experiments and edited the manuscript. F.W.S. performed ChIP-seq experiments. S.Pu. and U.B. performed computational data analysis for ChIP-seq. G.Z. generated stable shRNA-mediated knockdown cell lines.

## FUNDING

This work was supported by the Ontario Research Fund from the Ontario Ministry of Research and Innovation and by CIHR Operating Grant [PJT-148925] to J.F.G. D.Y.Z. was supported by a National Science and Engineering Research Council of Canada Studentship and an Ontario Graduate Scholarship. U.B. was supported by a long-term Postdoctoral Fellowship from HFSP. B.J.B. holds the University of Toronto Banbury Chair in Medical Research.

## CONFLICT OF INTEREST

The authors declare no competing interests.

