## Supplementary data file for "Regulation of transcription termination by FUS and TDP-43"

**Manuscript title:** Regulation of transcription termination by FUS and TDP-43

Table 1

| RNAPII 3'-end processing and termination factors | Detection method |
|---|---|
| CPSF5 | Rme2s Antibody IP-MS |
| CPSF6 (GR) | Rme2s Antibody IP-MS |
| CPSF1 | Rme2s Antibody IP-MS |
| CSTF2 (PRG, TRG) | SILAC labeling for MS |
| PABP1 (NR, LR, PR) | Rme2s Antibody IP-MS |
| PABP2 (PR, GR) | Rme2s Antibody IP-MS |
| PABP4 (PR) | SILAC labeling for MS |
| DHX9 (PRP, QRG, GRG) | SILAC labeling for MS |
| PCF11 (PR) | SILAC labeling for MS |
| WDR33 (GR) | SILAC labeling for MS |
| EF1a1/2 | SILAC labeling for MS |
| DDX20 | SILAC labeling for MS |
| RBM9 | SILAC labeling for MS |
| RBBP6 | Rme Antibody IP-MS |
| TAF15 (GR) | SILAC labeling for MS |
| G3BP1/2 (GR, PR) | SILAC labeling for MS |
| FCP1 (GR, PR) | Rme2s Antibody IP-MS |
| SPT5 (GR) | Rme2s Antibody IP-MS |
| EWSR1 (GR) | Rme2s Antibody IP-MS |
| FUS (GR) | Rme2s Antibody IP-MS |
| TDP-43 (GR) | Rme Antibody IP-MS |
| XRN2 (GR, PRG, YRP) | SILAC labeling for MS |

Table 2

| ALS Mutated Genes | Frequency | Proteins trapped in cytoplasmic inclusion bodies | |
|---|---|---|---|
| | | TDP-43 | FUS |
| C9ORF72 expansion | 40% | yes | |
| Superoxide Dismutase (SOD1) | 20% | | |
| Granulin(GRN) | low | yes | |
| Angiogenin, Ribonuclease (ANG) | low | yes | |
| Ubiquilin 2 (UBQLN2) | low | yes | |
| Profilin 1 (PFN1) | low | yes | |
| valosin containing protein (VCP) | low | yes | |
| hnRNPA1/B1 | low | yes | |
| Sequestosome 1 (SQSTM1) | low | yes | |
| Ataxin 2 (ATXN2) | low | yes | |
| Optineurin (OPTN) | low | yes | |
| TAR DNA binding protein (TDP-43) | low | yes | |
| Fused in Sarcoma (FUS) | low | | yes |
| TATA-Binding Protein-Associated Factor (TAF15) | low | | Yes |
| EWS RNA-binding protein 1 (EWSR1) | low | | yes |
| Senataxin (SETX) | low | | |

**Supplementary Table 1** RNAPII 3'-end processing, elongation, and termination factors that contain arginine dimethylations as potential SMN binders.

**Supplementary Table 2** Genes implicated in the pathology of ALS that often lead to the formation of cytoplasmic inclusion bodies that trap TDP-43 or FUS.

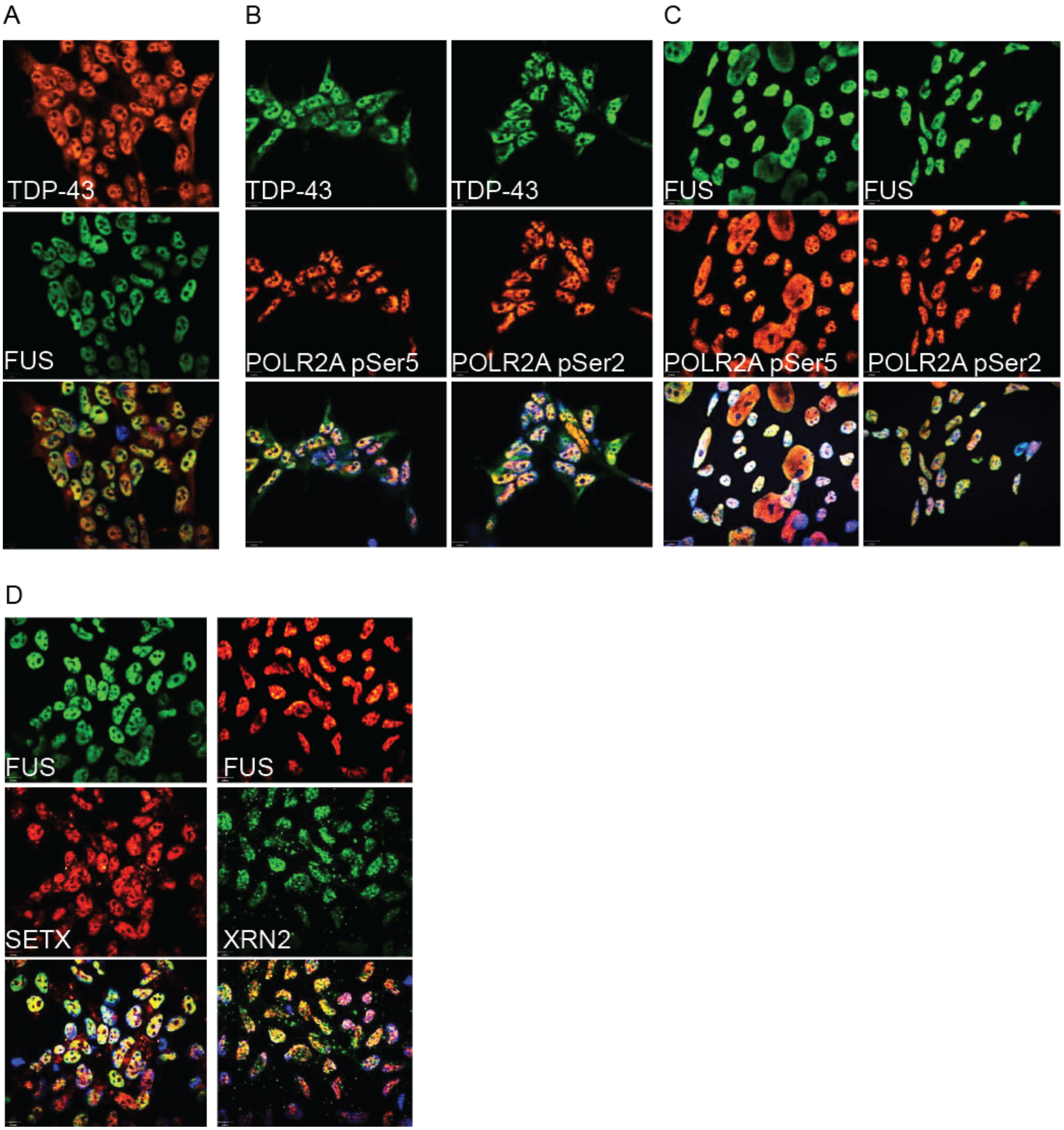


**Supplementary Figure S1** FUS and TDP-43 colocalize in nuclei with RNAPII

**A-D.** Immuno-staining of FUS and TDP-43 in HEK293 cells. The staining showed that FUS and TDP-43 colocalize with each other in the nuclei, with termination factors such as SETX and XRN2, and with POLR2A (pSer2, pSer5). Hoechst stain for DNA shown in blue.

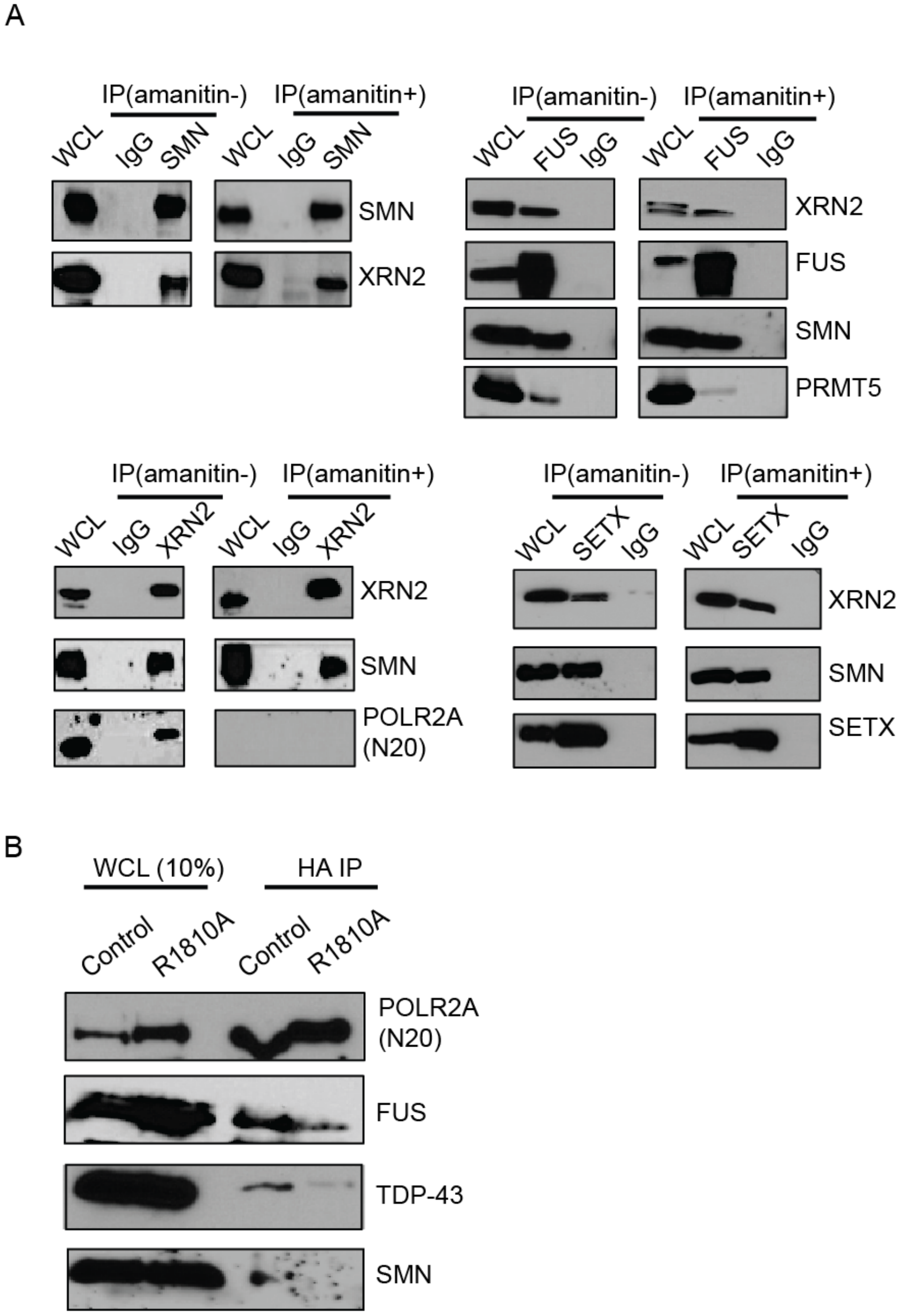


**Supplementary Figure S2** SMN and FUS interact with termination factors independently of RNAPII, and the POLR2A R1810A mutation disrupts the interactions amongst RNAPII, SMN, TDP-43, and FUS

**A.** IP with the indicated antibodies from HEK293 whole cell lysates (WCL), followed by western blotting with the indicated antibodies with or without α-amanitin pre-treatment of the HEK293 cells to abolish RNAPII. Many of the interactions occur independently of RNAPII, as they persist with α-amanitin treatment, including SMN-XRN2, SMN-FUS, SETX-XRN2, SETX-FUS, FUS-XRN2.

**B.** IP with the indicated antibodies from Raji WCL stably expressing the HA-tagged wild-type or R1810A mutant POLR2A upon 3-day α-amanitin treatment (2ug/ml) that abolishes the endogenous POLR2A. Anti-HA was used to precipitate HA-tagged wild-type (Control) or R1810A mutant POLR2A, followed by western blotting with the indicated antibodies.

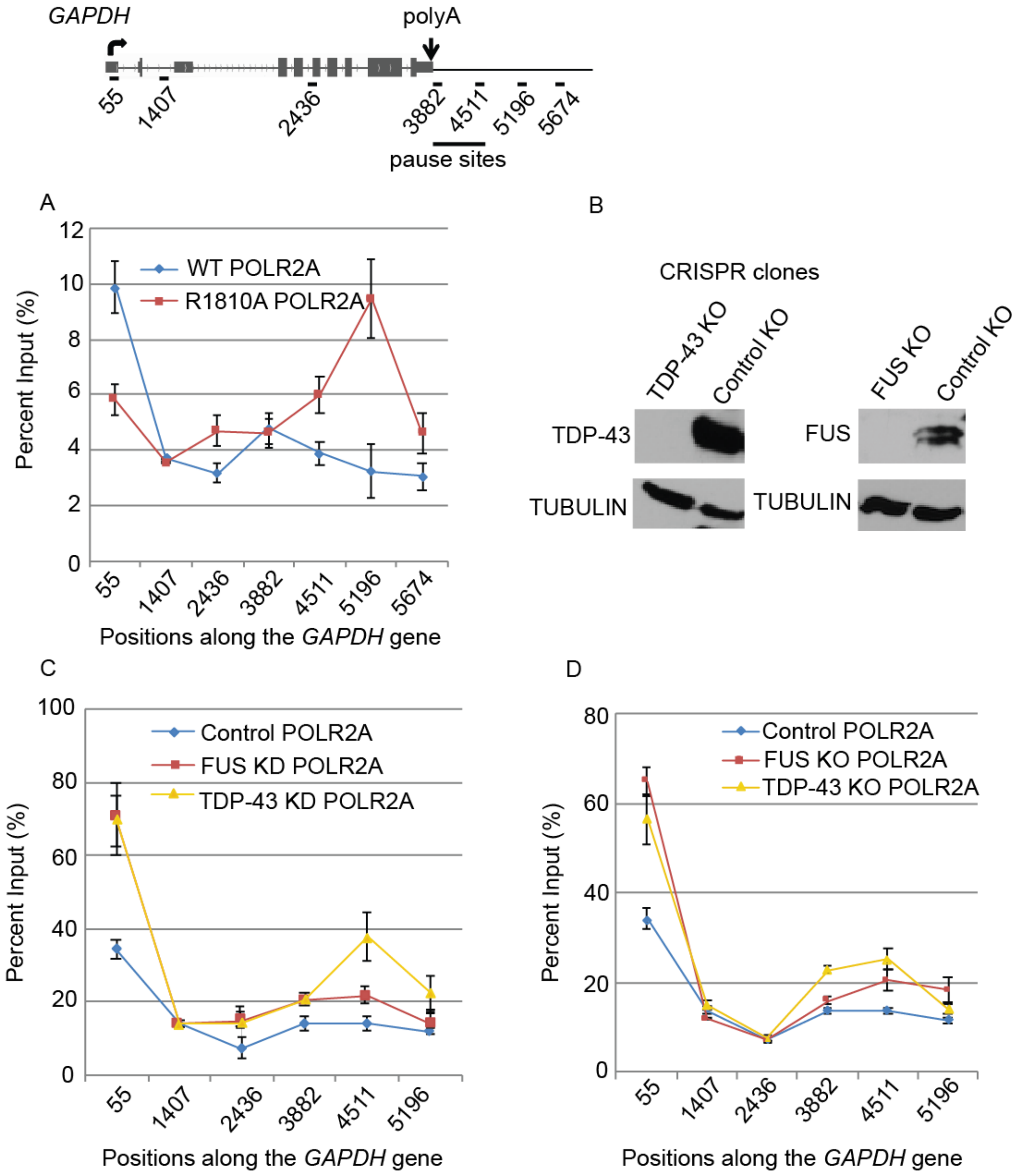


**Supplementary Figure S3** RNAPII R1810, FUS and TDP-43 regulate RNAPII termination on the *GAPDH* gene

**A.** ChIP quantification of WT and the endogenous R1810A mutant RNAPII using POLR2A antibody (8WG16, N20) in HEK293 cells, using the indicated primer positions for qPCR along the *GAPDH* gene. Error bars denote biological replicates as s.e.m. (n= 4).

**B.** Western blotting with the indicated antibodies verifies the knock-out of TDP-43 or FUS by the CRISPR/Cas9 system.

**C.** Quantification of RNAPII ChIP using POLR2A antibody (4H8, N20) in HEK293 cells, using the indicated primer positions for qPCR along the *GAPDH* gene, after stably knocking down FUS or TDP-43, with GFP knock-down as a negative control. Error bars denote biological replicates as s.e.m. (n= 3).

**D.** Quantification of RNAPII ChIP using POLR2A antibody (4H8, N20) in HEK293 cells, using the indicated primer positions for qPCR along the *GAPDH* gene, after knocking out FUS or TDP-43 using the CRISPR/Cas9 system or using scrambled guide RNA as negative control. Error bars denote biological replicates as s.e.m. (n = 3).

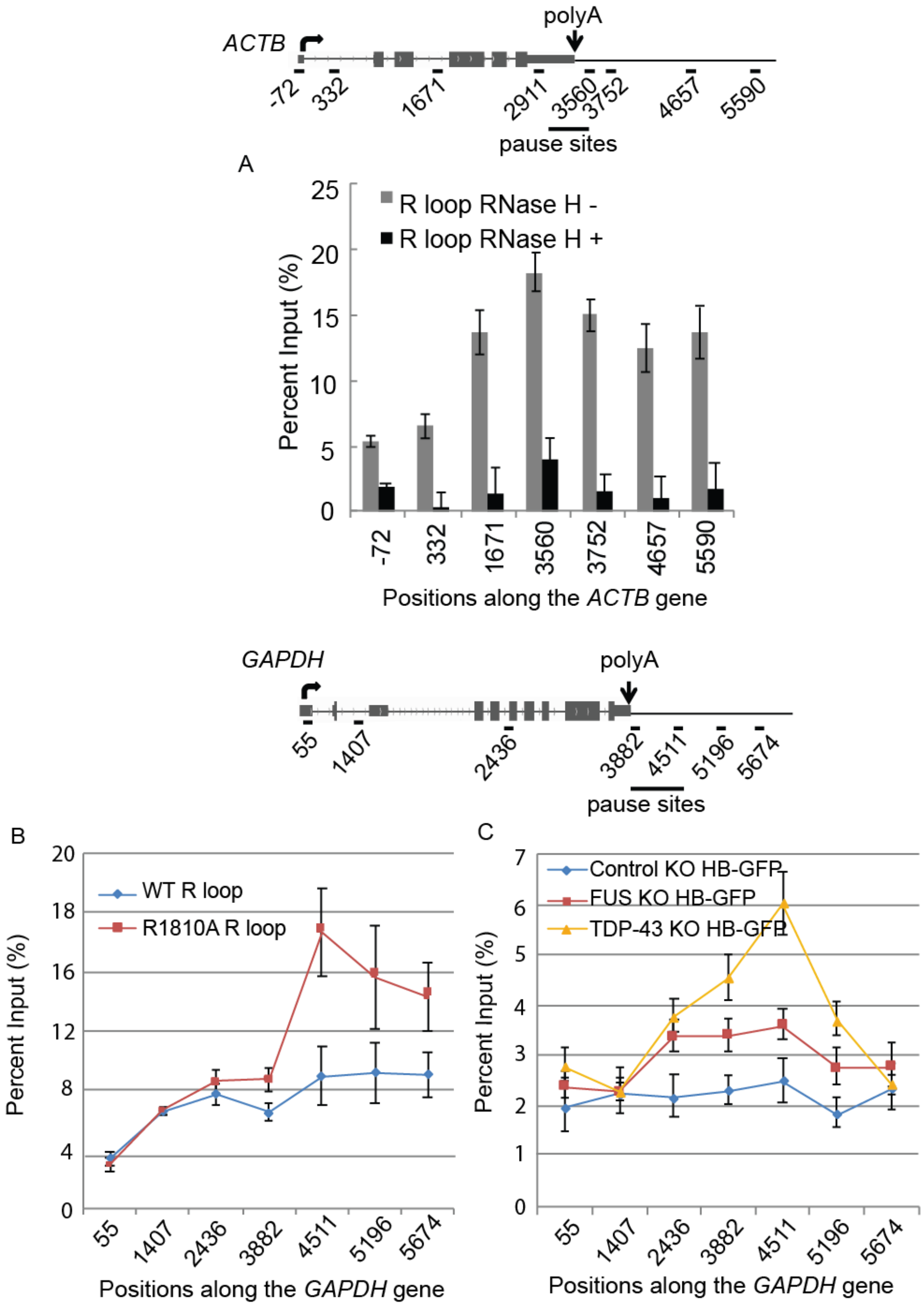


**Supplementary Figure S4** RNAPII R1810, FUS, and TDP-43 are important for resolving R-loops in RNAPII termination regions

**A.** Quantification of R-loops with the S9.6 antibody in HEK293 cells with or without RNase H treatment on the *ACTB* gene. This control experiment was previously published (9).

**B.** Quantification of R-loops with the S9.6 antibody for the WT and the endogenous R1810A mutant RNAPII in HEK293 cells, with the indicated primer positions for qPCR along the *GAPDH* gene. Error bars denote biological replicates as s.e.m. (n=3).

**C.** Quantification of R-loops with the GFP-HB construct, with the indicated primer positions for qPCR along the *GAPDH* gene, after knocking out FUS or TDP-43 with the CRISPR/Cas9 system or with the scrambled guide RNA as a negative control. Error bars denote biological replicates as s.e.m (n= 3).

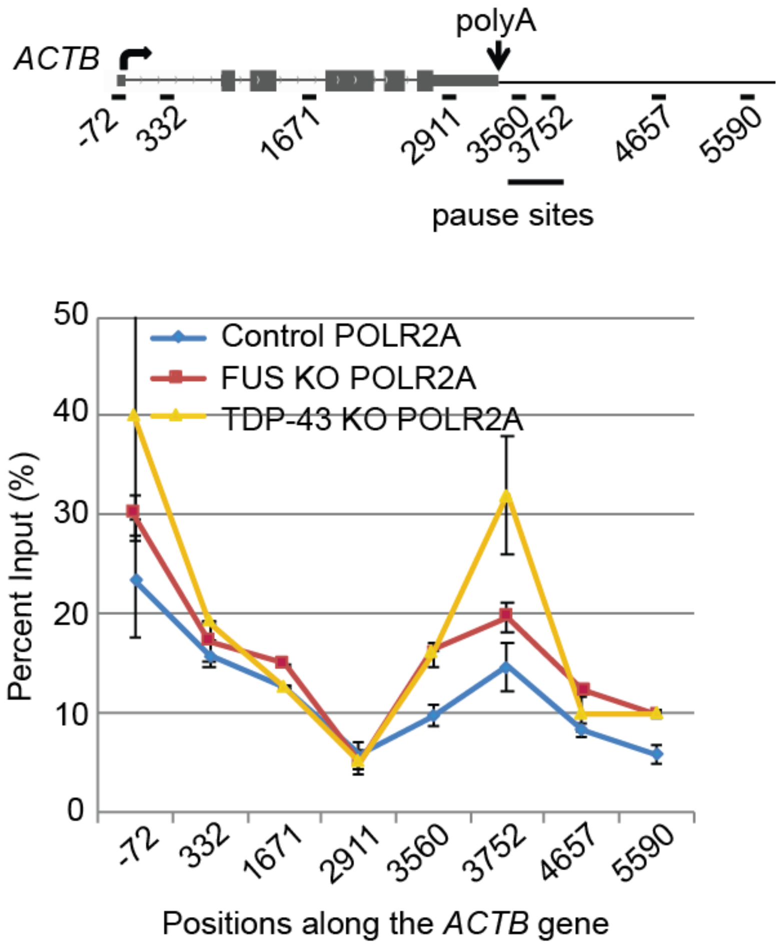


**Supplementary Figure S5** FUS and TDP-43 regulate RNAPII termination on the *ACTB* gene

Quantification of RNAPII ChIP using POLR2A antibody (4H8, N20) in HEK293 cells, using the indicated primer positions for qPCR along the *ACTB* gene after knocking out FUS or TDP-43 with the CRISPR/Cas9 system, or using scrambled guide RNA as negative control.  Error bars denote biological replicates as s.e.m. (n= 5).

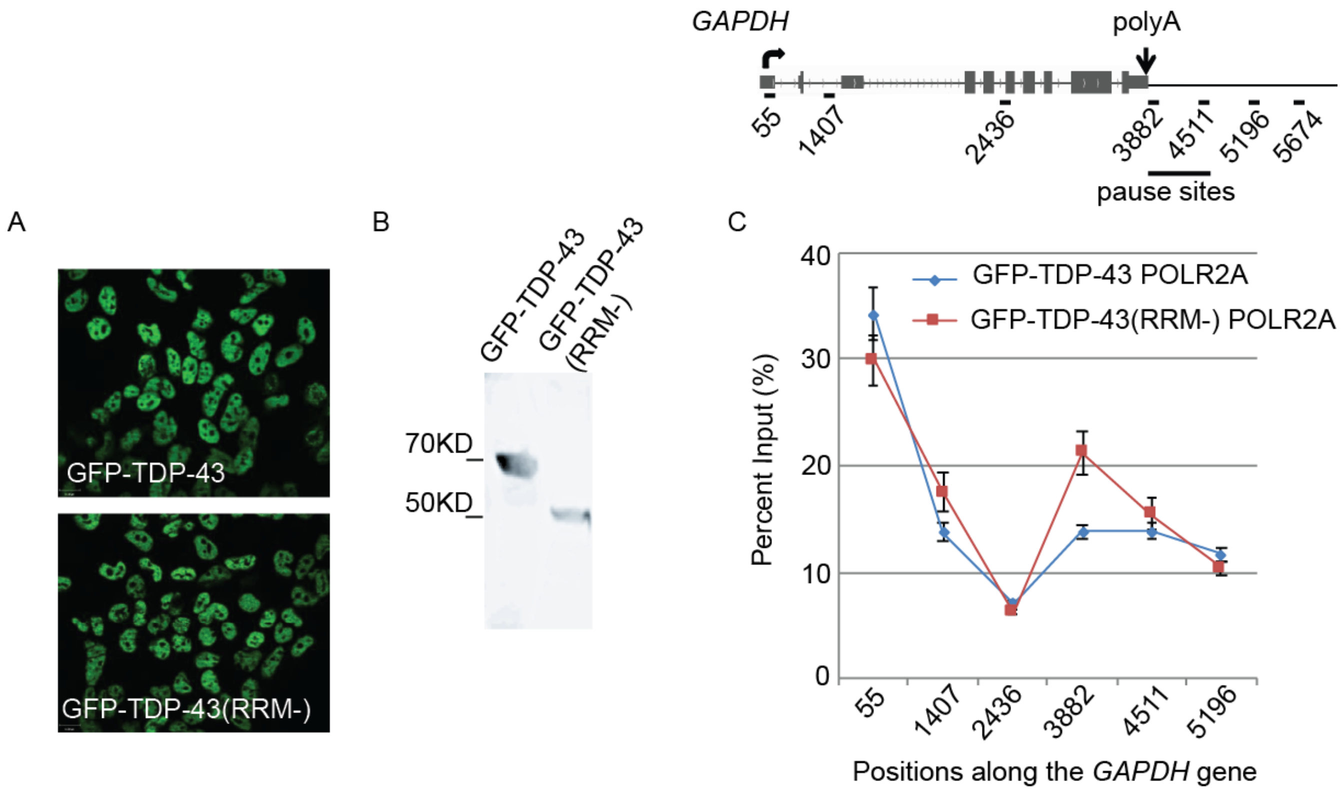


**Supplementary Figure S6** RRM domains of TDP-43 are important for TDP-43 to regulate RNAPII termination on the *GAPDH* gene

**A.** Immuno-staining verifies the expression pattern of full length and the truncated GFP-TDP-43 (RRM-) induced upon 1 day of doxycycline treatment.
**B.** Western blotting with GFP antibody to show the levels and sizes of the GFP fusion proteins extracted from the HEK293 WCL. The truncated TDP-43 lacks RRM (RNA recognition motifs) at the C terminus of the protein.
**C.** Quantification of RNAPII ChIP using the POLR2A antibody (N20) with cells overexpressing wild type GFP-TDP-43 or the GFP-TDP-43 (RRM-) truncation mutant, using the indicated primer positions for qPCR along *GAPDH* genes. Error bars denote biological replicates as s.e.m. (n= 3).

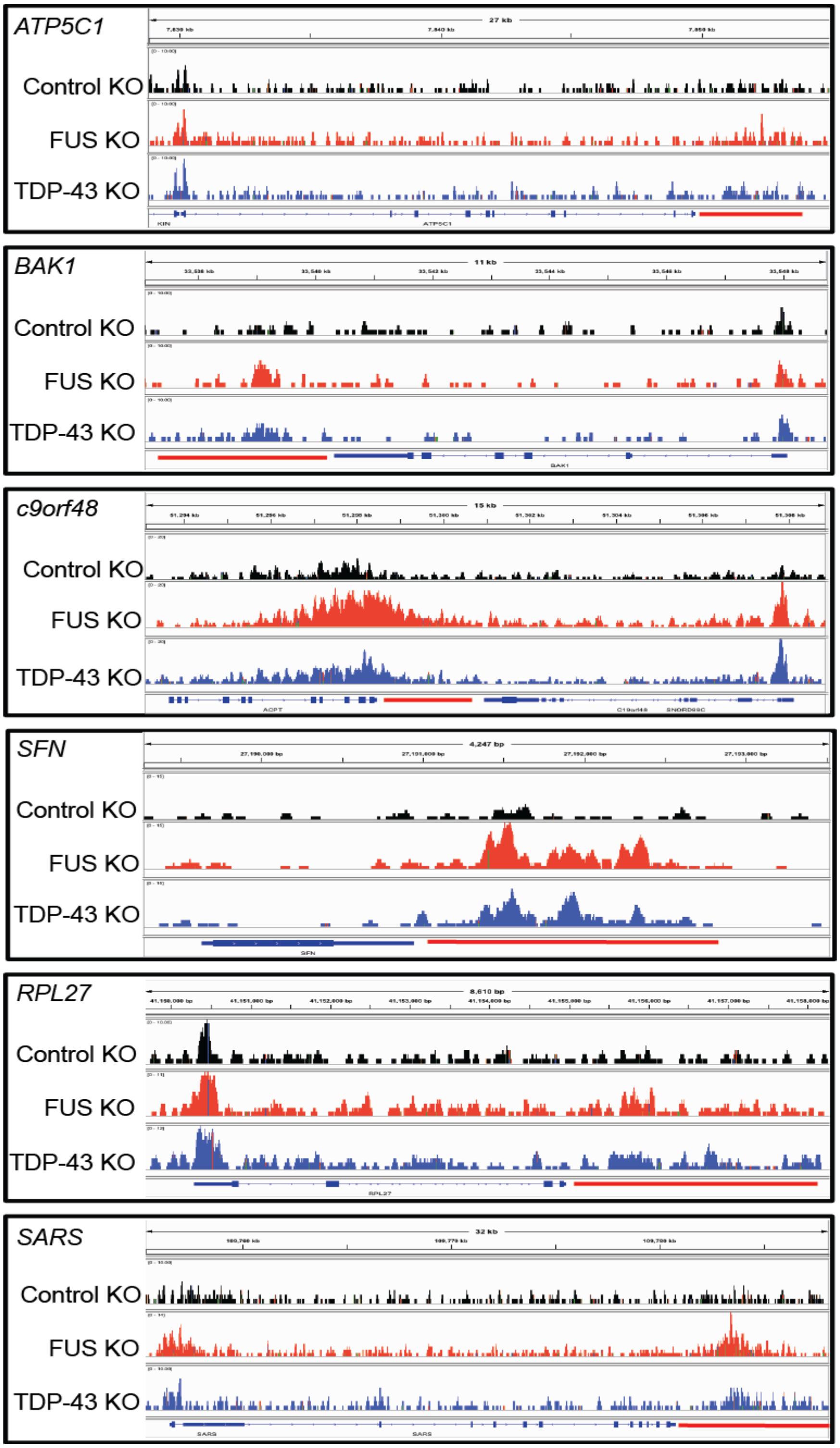


**Supplementary Figure S7** FUS and TDP-43 regulate RNAPII promoter and terminator stalling

RNAPII ChIPseq results for several housekeeping genes are displayed in detail with the Integrative Genomics Viewer. RNAPII termination regions are underlined in red.

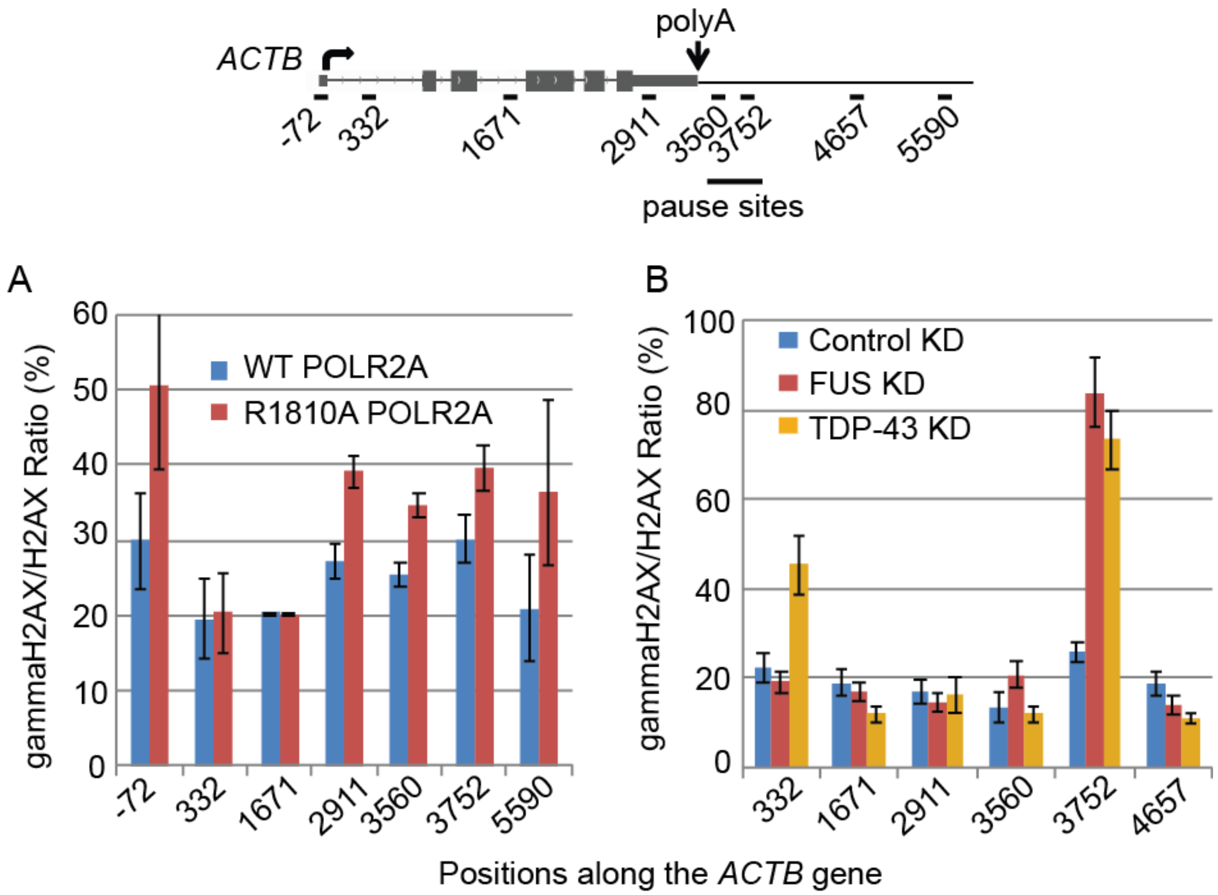


**Supplementary Figure S8** RNAPII CTD R1810, FUS and TDP-43 prevent DNA damage at RNAPII transcription terminators

**A, B.** ChIP quantifications of the γH2AX/H2AX ratio in HEK293 cells along the length of the *ACTB* gene, comparing WT and the endogenous R1810A mutant POLR2A (**A**), or after knocking down FUS or TDP-43, with the knock-down of GFP as a negative control (**B**).  Error bars denote biological replicates as s.e.m. (n=4).
